# The organosulfur compound dimethylsulfoniopropionate (DMSP) is utilized as an osmoprotectant by *Vibrio* species

**DOI:** 10.1101/2020.09.10.292482

**Authors:** Gwendolyn J. Gregory, Katherine E. Boas, E. Fidelma Boyd

**Affiliations:** Department of Biological Sciences, University of Delaware, Newark, DE, 19716

## Abstract

Dimethylsulfoniopropionate (DMSP) is a key component of the global geochemical sulfur cycle that is a secondary metabolite produced in large quantities by marine phytoplankton and utilized as an osmoprotectant. Bacterial DMSP lyases convert DMSP to the climate active gas dimethylsulfide (DMS). Whether marine bacteria can also accumulate DMSP as an osmoprotectant to maintain the turgor pressure of the cell in response to changes in external osmolarity remains unknown. The marine halophile *Vibrio parahaemolyticus*, contains at least six osmolyte transporters, four betaine carnitine choline transport (BCCT) carriers BccT1-BccT4 and two ABC-family ProU transporters. In this study, we showed that DMSP is used as an osmoprotectant by *V. parahaemolyticus* and several other *Vibrio* species including *V. cholerae* and *V. vulnificus*. Using a *V. parahaemolyticus proU* double mutant, we demonstrated that these ABC transporters are not required for DMSP uptake. However, a *bccT* null mutant lacking all four BCCTs had a growth defect compared to wild type in high salt media supplemented with DMSP. Using *bccT* triple mutants, possessing only one functional BCCT, in growth pattern assays, we identified two BCCT-family transporters, BccT1 and BccT2 are carriers of DMSP. *Vibrio cholerae* and *V. vulnificus*, only contain a homolog of BccT3 and functional complementation in *Escherichia coli* MKH13 showed only *V. cholerae* BccT3 could transport DMSP. In *V. vulnificus* strains, we identified and characterized an additional BCCT transporter that was also a carrier for DMSP. Phylogenetic analysis uncovered at least 11 distinct BCCT transporters among members of the Harveyi clade, with some species having up to 9 BCCTs as exemplified by *V. jasicida*.

**Importance:** DMSP is present in the marine environment, produced in large quantities by marine phytoplankton as an osmoprotectant, and is an important component of the global geosulfur cycle. The bacterial family *Vibrionaceae* is comprised of marine species, many of which are halophiles such as *V. parahaemolyticus*, which can utilize a wide range of osmolytes and possesses at least six transporters for the uptake of these compounds. Here, we demonstrated that *V. parahaemolyticus* and other *Vibrio* species can accumulate DMSP as an osmoprotectant and show that the BCCT family transporters were required. DMSP was transported by four different BCCT transporters; BccT1, BccT2, BccT3 and BccT5 depending on the species. Bioinformatics and phylogenetics demonstrated that *Vibrio* species contain a large number of BCCTs and that many of these are associated with different metabolic pathways.

## Introduction

Compatible solutes (osmolytes) are accumulated by bacteria to maintain the turgor pressure of the cell in response to high external osmolarity (1-3). Known compatible solutes of bacteria include glycine betaine, ectoine, proline, glutamate, glycerol and trehalose (2, 4-9). Compatible solutes, as the name suggests, are compounds that can be accumulated to high levels and are compatible with the molecular machinery and processes of the cell. These osmolytes allow organisms to continue to grow and divide in unfavorable environments that may be characterized as changes in osmolarity and temperature shifts (10-19). The uptake and biosynthesis of compatible solutes in response to osmotic stress has been studied extensively in *Escherichia coli* and *Bacillus subtilis*, both can biosynthesize glycine betaine from choline and can uptake glycine betaine and choline amongst other osmolytes (10, 20-32).

Organisms that live in marine environments have adapted to grow optimally in high salinity habitats and also have evolved mechanism to maintain cellular homeostasis to cope with fluctuations in salinity that they encounter. Members of the family *Vibrionaceae* have the ability to grow optimally at 0.5 M-1.0 M NaCl, but regularly encounter fluctuations in osmolarity ranging from 0.1 M to 1.5 M NaCl (33). One of the key responses of *Vibrio* species to fluctuations in osmolarity is the accumulation of compatible solutes, either through biosynthesis of the osmolytes ectoine and glycine betaine or the rapid uptake of these osmolytes from the environment using as many as six osmolyte transporters (33-35). *Vibrio parahaemolyticus*, a halophile ubiquitous in the marine environment, biosynthesizes glycine betaine from exogenously supplied choline, and ectoine from aspartic acid *de novo* and can uptake at least 14 different compatible solutes (33-36). This species contains two osmolyte ATP Binding Cassette (ABC)-family transporters, ProU1 (VP1726-VP1729) and ProU2 (VPA1112-VPA1114), and four Betaine Carnitine Choline Transporter (BCCT) family transporters, BccT1 (VP1456), BccT2 (VP1723), BccT3 (VP1905), and BccT4 (VPA0356) (33). All BCCTs described have 12 predicted *trans*-membrane (TM) α-helical domains, TM1 to TM12, which is a defining feature in their classification (21, 37-41). In addition to the 12 TM domains, these proteins contain hydrophilic N- and C-terminal tails of varying length (21, 39-41). Some species such as *V. vulnificus, V. cholerae* and *Aliivibrio fischeri* contain only a homolog of BccT3 (33). We have previously demonstrated that BCCTs can transport glycine betaine, choline, dimethylglycine (DMG), ectoine, and proline, with some redundancy in the substrates transported by each BCCT (35, 36).

Dimethylsulfoniopropionate (DMSP) is an organosulfur compound abundant in marine surface waters, produced by phytoplankton and some halophytic vascular plants in large quantities and used by these primary producers as an osmoprotectant, thermoprotectant and antioxidant (42-47). DMSP is an important component of the global geochemical sulfur cycle as a precursor for dimethylsulfide (DMS). DMS is a climate active gas that is produced from the degradation of oceanic DMSP, releasing sulfur-containing aerosols into the atmosphere (44, 45, 48-52). It was reported that DMSP is also present in marine and estuarine sediments, and is produced by bacteria in these environments (53, 54). Phytoplankton blooms that have high production of DMSP are sources of reduced sulfur and carbon for many marine heterotrophic bacteria (43, 55-59). DMSP-catabolizing bacteria are mainly confined to marine alpha-proteobacteria, SAR11, SAR116 and Roseobacter (60-67). Given that many species of *Vibrionaceae* interact and associate with DMSP producers, surprisingly there are no studies on the use of DMSP as an osmoprotectant in these bacteria, or in marine bacteria in general (3, 68-70). The first direct evidence that DMSP can be used as an osmoprotectant for bacteria came from a study in *E. coli*, which does not encounter this compound naturally. Osmotically stressed *E. coli* responded to DMSP and the ABC-family ProU transporter was shown to uptake DMSP to relieve salt stress (68). A more recent study in the Gram-positive soil bacterium *B. subtilis* showed that DMSP was an osmoprotectant transported into the cell by the ABC-family OpuC transporter (69). The BCCT family transporter DddT linked to DMSP cleavage pathways in *Halomonas* sp HTNK1 was demonstrated to uptake DMSP in a heterologous *E. coli* background (67, 70). Outside of these studies, little is known about bacterial utilization of DMSP as an osmolyte or the bacterial response to this abundant marine compound.

In this study, we examined *Vibrio* species for their ability to utilize DMSP as an osmoprotectant. We determined whether a *V. parahaemolyticus ectB* mutant, which cannot grow in high NaCl in the absence of an exogenous osmolyte, could be rescued in M9 minimal media glucose (M9G) supplemented with DMSP. Then we investigated the effectiveness of DMSP as an osmoprotectant in several *Vibrio* species including *V. cholerae* and *V. vulnificus*, to determine whether this was a general trait of this genus. *Vibrio parahaemolyticus* has six osmolyte transporters, two ProUs and four BCCTs. First, we examined whether a *proU* double mutant had a defect when grown in high salt with DMSP as an osmolyte. Then, we examined the role of each of the BCCTs in DMSP uptake using four triple mutants that each contained a single *bccT* gene. *Vibrio cholerae* contains only a single BCCT family transporter, a BccT3 homolog, and we determined its ability to uptake DMSP. In *V. vulnificus*, a homolog of BccT3 is present and we identified an additional BCCT family transporter named BccT5 that could uptake DMSP. Phylogenetic analysis showed BCCT transporters are prevalent among members of the Harveyi clade and can contain up to 9 divergent BCCT transporters.

## Results

### *Vibrio parahaemolyticus* can utilize dimethylsulfoniopropionate (DMSP) as a compatible solute

DMSP is produced in large quantities by phytoplankton and used by these primary producers as an osmolyte. However, the role of DMSP as an osmolyte for marine bacteria remains largely unknown. As a halophile, *V. parahaemolyticus* grows optimally in 500 mM NaCl (∼3% NaCl), and can grow in salinities up to 1.5 M (∼9%) (34). We examined the ability of *V. parahaemolyticus* to grow in high salinity (6% NaCl) conditions using DMSP as an osmolyte. The *V. parahaemolyticus ectB* deletion mutant that cannot grow in high NaCl growth conditions in the absence of exogenous osmolytes, as was previously shown was used as a positive control (34, 36). We examined growth of wild type and the *ectB* mutant in M9 minimal media supplemented with glucose and 6% NaCl (M9G 6%NaCl), in the presence and absence of DMSP. Growth pattern analysis after 24 h showed that the lag phase of the wild-type strain without exogenous compatible solutes is approximately six hours with a growth rate of 0.042 h^-1^, while the Δ*ectB* mutant did not grow (36) (**Fig. 1**). However, in M9G 6% NaCl supplemented with DMSP the *ectB* mutant grew similarly to wild type, with growth rates of 0.082 h^-1^ and 0. 074 h^-1^, respectively. Lag phases were reduced to ∼3 h for the wild-type strain and <1 h for the *ectB* mutant. This demonstrated that *V. parahaemolyticus* can uptake and utilize DMSP as a highly effective osmolyte (**Fig. 1**).

**Figure 1.**
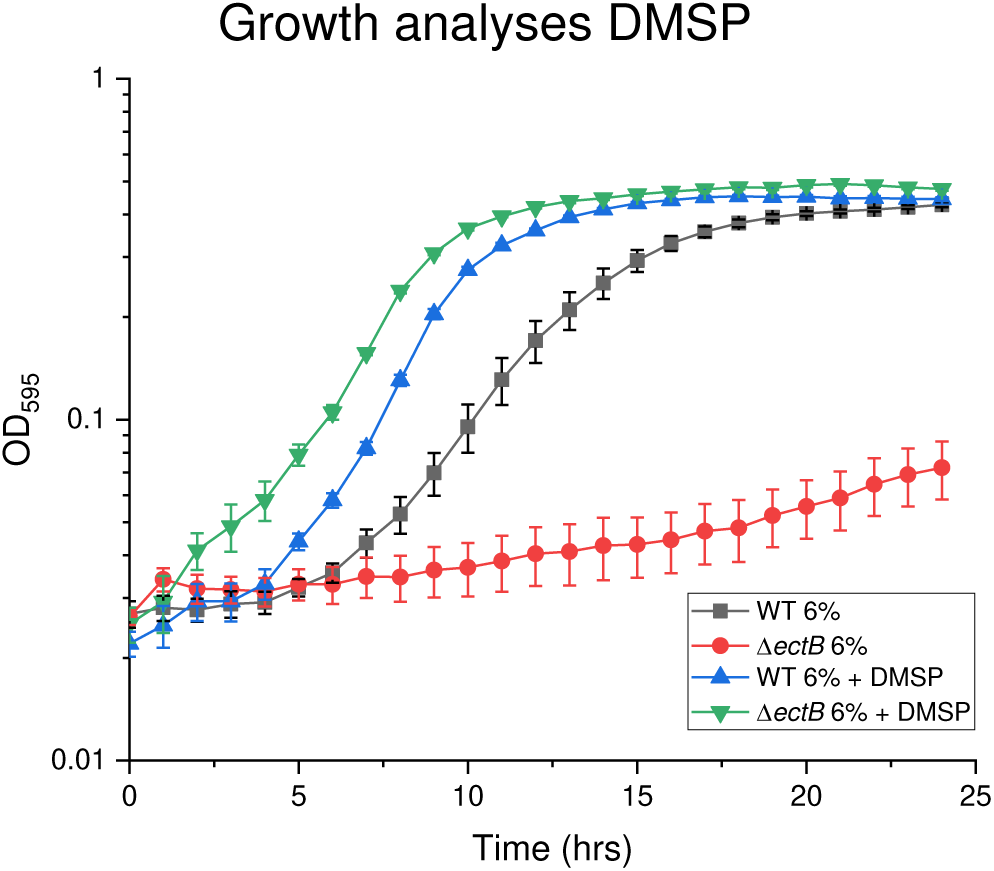
Growth analysis of wild-type (WT) *V. parahaemolyticus* RIMD2210633 and an *ectB* mutant was conducted in M9G supplemented with 6% NaCl and DMSP. Optical density (OD_595_) was measured every hour for 24 hours; mean and standard error of at least two biological replicates are displayed.

### DMSP is utilized as an osmolyte by *Vibrio* species

To determine whether other *Vibrio* species can utilize DMSP as an osmolyte, we tested DMSP uptake in *V. harveyi* 393, *V. fluvialis* ATCC33809, *V. vulnificus* YJ016, and *V. cholerae* N16961. The growth of *V. harveyi* 393, *V. vulnificus* YJ016, and *V. cholerae* N16961 was examined in M9G 4% NaCl and *V. fluvialis* ATCC33809 in M9G 5% NaCl, with and without added DMSP (**Fig. 2A-D**). The growth of *V. harveyi* 393 showed a 10 h lag phase with a growth rate of 0.06 h^-1^, however, the addition of DMSP resulted in a lag phase of less than 2 h, with a maximum growth rate of 0.088 h^-1^. *Vibrio vulnificus* did not grow in M9G 4% NaCl. However, the addition of DMSP rescued growth with a growth rate of 0.065 h^-1^, but cells had a 13h lag phase suggesting uptake was not efficient or that DMSP is not a very effective compatible solute for *V. vulnificus* (**Fig. 2B**). *V. fluvialis* had a growth rate of 0.026 h^-1^ with a lag phase of ∼4 h when grown without DMSP. The lag phase was reduced to ∼2 h and the growth rate increased to 0.041 h^-1^ in the presence of DMSP (**Fig. 2C**). We found that *V. cholerae* cells had a reduced lag phase, from 2 h to less than 1 h, and an increased growth rate, from 0.034 h^-1^ to 0.059 h^-1^, in the presence of DMSP (**Fig. 2D**).These data demonstrated that all *Vibrio* species tested can utilize DMSP as an osmolyte.

**Figure 2.**
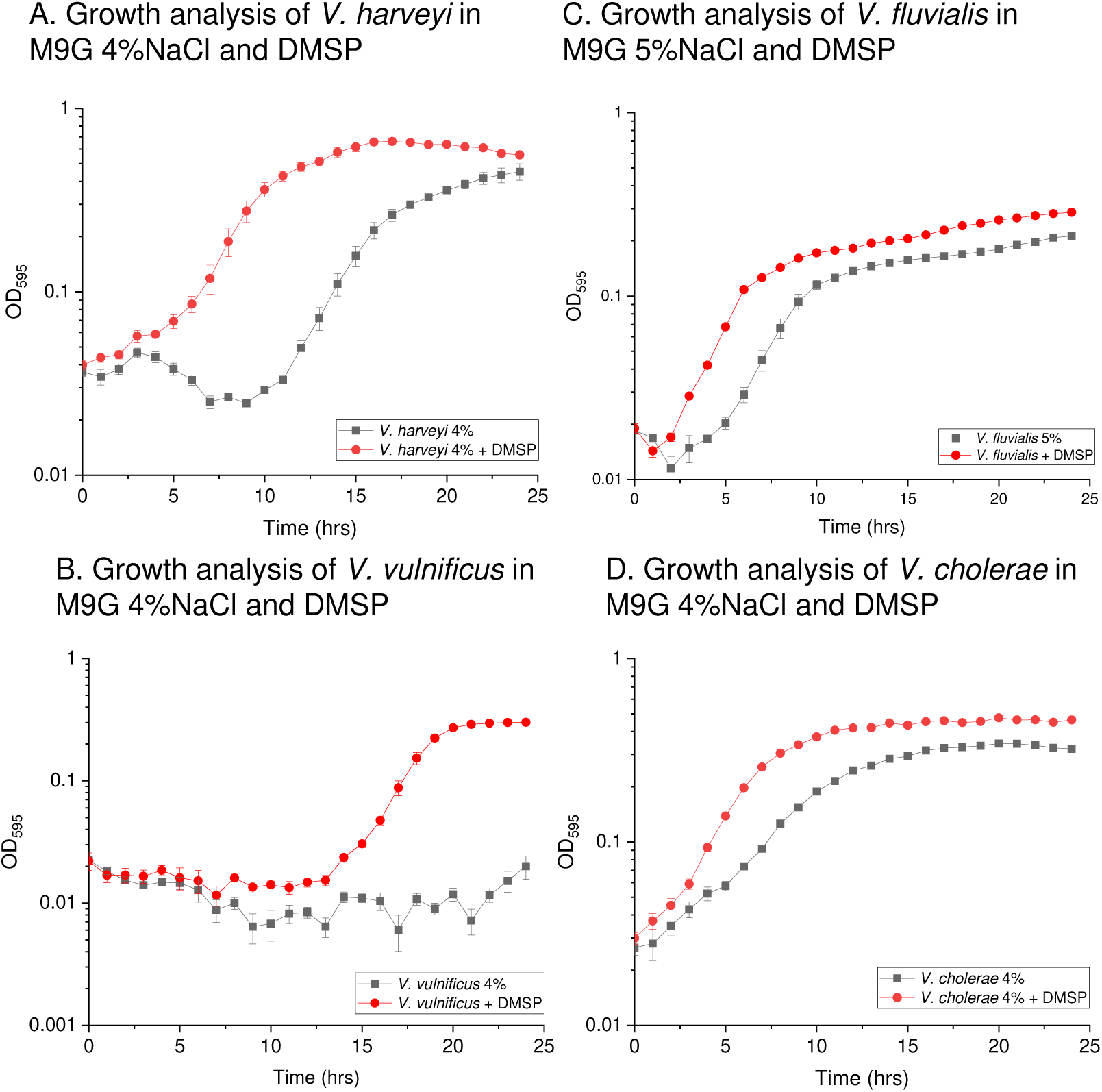
Growth analyses of **(A)** *V. harveyi* 393, **(C)** *V. vulnificus* YJ016, and **(D)** *V. cholerae* N16961 were conducted in M9G 4% NaCl with and without exogenous DMSP. Growth analyses of **(B)** *V. fluvialis* ATCC was conducted in M9G 5% NaCl with and without exogenous DMSP. Optical density (OD_595_) was measured every hour for 24 hours. Mean and standard error of two biological replicates are shown.

Some marine bacteria are able to catabolize DMSP (71-74). To ensure that the reduced lag phase in each *Vibrio* species tested was not due to use of DMSP as a carbon source, we grew each strain in M9 minimal media with DMSP as the sole carbon source, utilizing M9G (glucose as the sole carbon source) as a control. None of the *Vibrio* species tested grew with DMSP as the sole carbon source and all grew on M9G, which indicated that they cannot catabolize DMSP (**Fig. S1**). This data demonstrated that DMSP is a *bona fide* compatible solute for the *Vibrio* species tested.

### BCCT transporters required for DMSP uptake

Previous studies in other species showed that DMSP was transported into cells by ABC-family transporters (68, 69). *V. parahaemolyticus* contains two ABC-type osmolyte transporters, ProU1 (VP1726-VP1729) and ProU2 (VPA1112-VPA1114). Growth analyses with a double *proU* mutant (Δ*proU1*Δ*proU2*) were performed to determine whether either is required for DMSP uptake. In these assays, no difference between the mutant and wild type in the absence or presence of DMSP was seen (**Fig. S2**). These data demonstrated that ProU transporters were not required for uptake of DMSP in *V. parahaemolyticus*. Next, we examined whether the four BccTs in *V. parahaemolyticus* were responsible for the uptake of DMSP into the cell. To accomplish this, wild type and a *bccT* null strain were grown in the presence or absence of DMSP in M9G 6% NaCl. The lag phase of the wild-type strain was reduced from ∼6 hours to ∼3 hours when grown in the presence of DMSP, which indicates that DMSP is a highly effective osmoprotectant for *V. parahaemolyticus* (**Fig. 3**). However, the *bccT* null mutant did not exhibit a reduced lag phase in the presence of DMSP (**Fig. 3**). This indicated that at least one of the BCCTs is responsible for transport of DMSP into the cell and confirms that neither ProU plays a role in uptake of DMSP.

**Figure 3.**
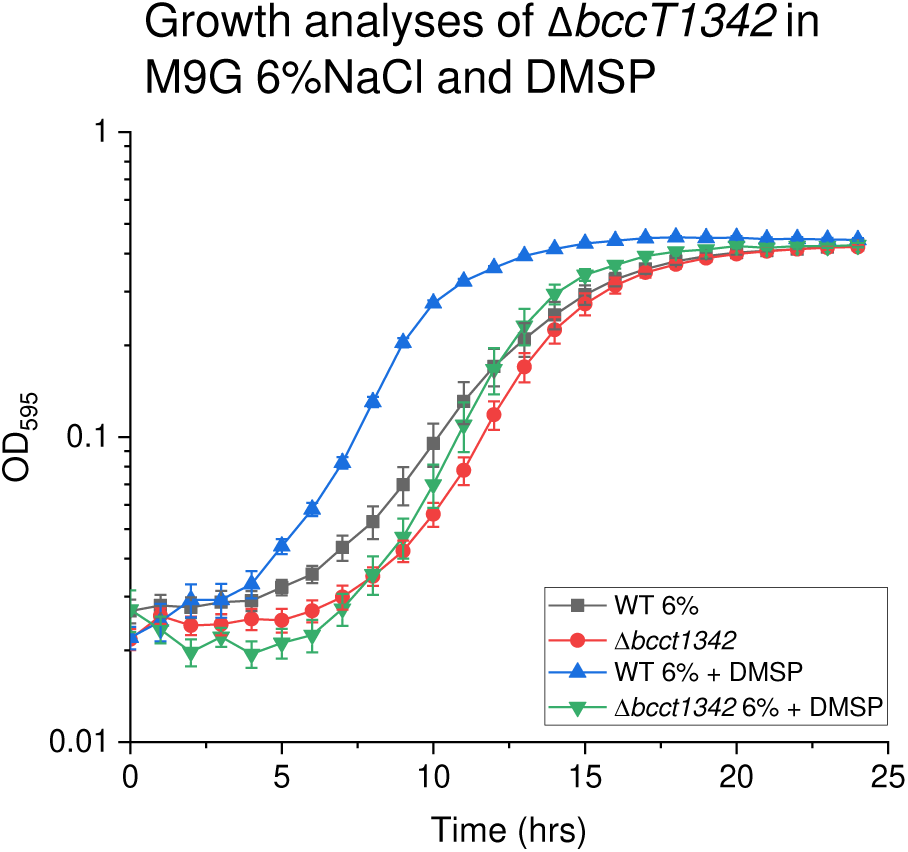
Growth analysis of wild-type (WT) *V. parahaemolyticus* RIMD2210633 and a Δ*bccT1*Δ*bccT3*Δ*bccT4*Δ*bccT2* mutant was conducted in M9G supplemented with 6% NaCl and DMSP. Optical density (OD_595_) was measured every hour for 24 hours; mean and standard error of at least two biological replicates are displayed.

To determine which of the BCCTs were responsible for uptake of DMSP, four triple *bccT* deletion mutants were utilized, each containing only one functional *bccT* gene. The Δ*bccT2*Δ*bccT3*Δ*bccT4* mutant, which expresses only *bccT1* (VP1456), had a slight reduction in lag phase and had a faster growth rate through the exponential phase when grown in the presence of DMSP. This suggests that BccT1 can transport DMSP with low efficiency (**Fig. 4A**). The Δ*bccT1*Δ*bccT3*Δ*bccT4* mutant, which expresses only *bccT2* (VP1723), showed a nearly identical reduction in lag phase as the wild-type strain, which indicated that it also transports DMSP into the cell (**Fig. 4B**). The Δ*bccT1*Δ*bccT2*Δ*bccT4* mutant, which expresses only *bccT3* (VP1905), and Δ*bccT1*Δ*bccT2*Δ*bccT3*, which expresses only *bccT4* (VPA0356), had slight reductions in their lag phase in the presence of DMSP (**Fig. 4C & 4D**).

**Figure 4.**
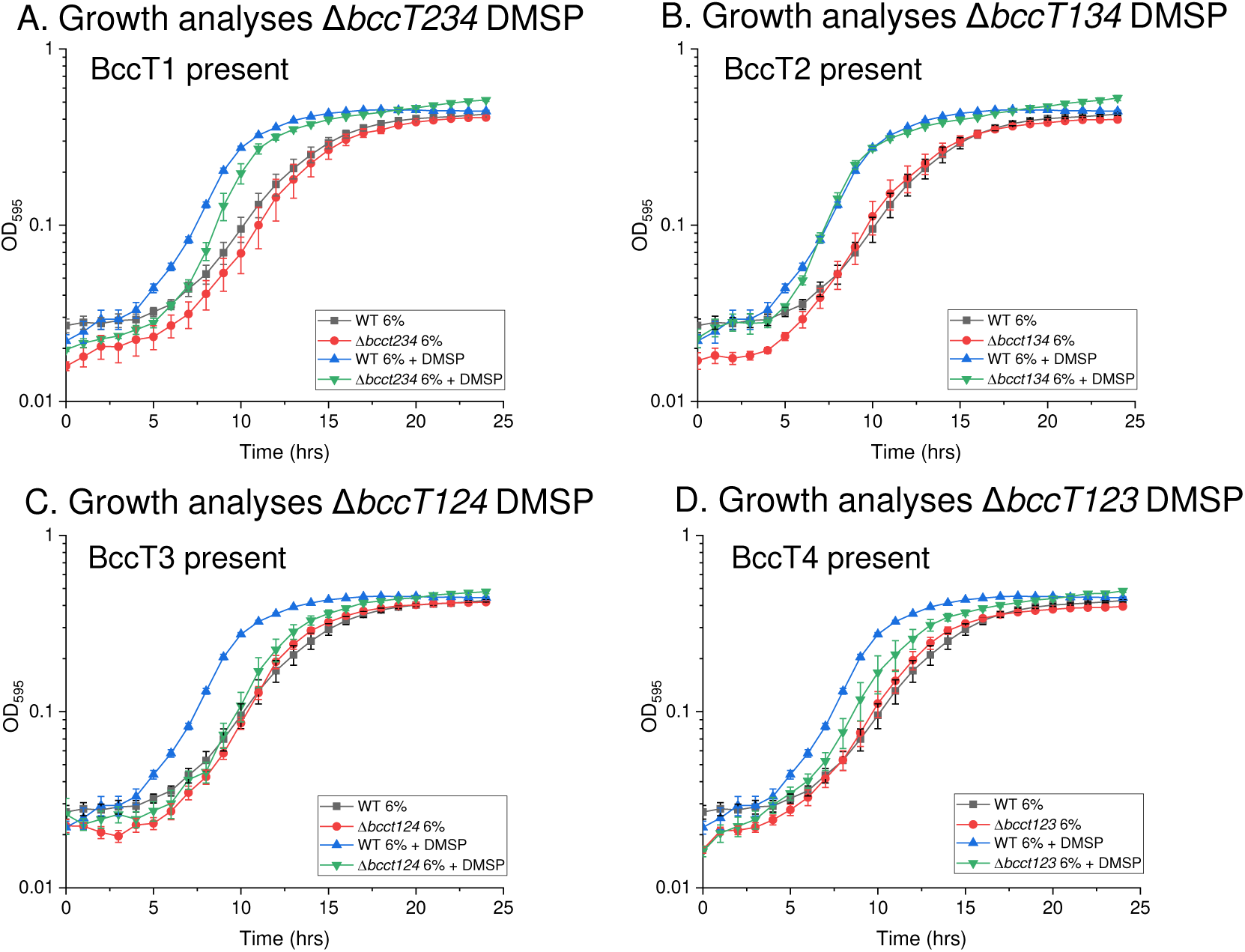
Growth analysis of wild-type (WT) *V. parahaemolyticus* RIMD2210633 and **(A)** Δ*bccT2*Δ*bccT3*Δ*bccT4*, **(B)** Δ*bccT1*Δ*bccT3*Δ*bccT4*, **(C)** Δ*bccT1*Δ*bccT2*Δ*bccT4*, or **(D)** Δ*bccT1*Δ*bccT2*Δ*bccT3* in M9G6% with and without the addition of exogenous DMSP. Optical density (OD_595_) was measured every hour for 24 hours; mean and standard error of at least two biological replicates are displayed.

Next, we examined DMSP transport capabilities of each BccT using functional complementation in *E. coli* strain MKH13, a mutant strain that has deletions in *betIBA-betT, proU, proP* and *putP* and cannot grow in M9G 4% NaCl (75). Strains were grown in M9G with 4% NaCl in the presence of DMSP, with an arabinose-inducible expression plasmid, pBAD33, harboring a full-length copy of a single *bccT*. We measured the optical density (OD_595_) after 24 hours for each complemented strain and compared to the empty vector strain. Only the BccT2-complemented strain (pBAVP1723) could grow in the presence of DMSP (**Fig. 5A**). The BccT1-complemented strain (pBAVP1456) was unable to grow in the presence of DMSP (**Fig. 5A**), although the *V. parahaemolyticus* strain containing only BccT1 transported DMSP, albeit with low efficiency (**Fig. 4A**). To confirm that *bccT1* is expressed and produces a functional protein in *E. coli*, we examined the ability of BccT1 to uptake glycine betaine, a known substrate for this transporter. In this assay, *E. coli* MKH13 complemented with *bccT1* (pBAVP1456) grew with the addition of glycine betaine, which indicated BccT1 is functional (**Fig. 5B**).

**Figure 5.**
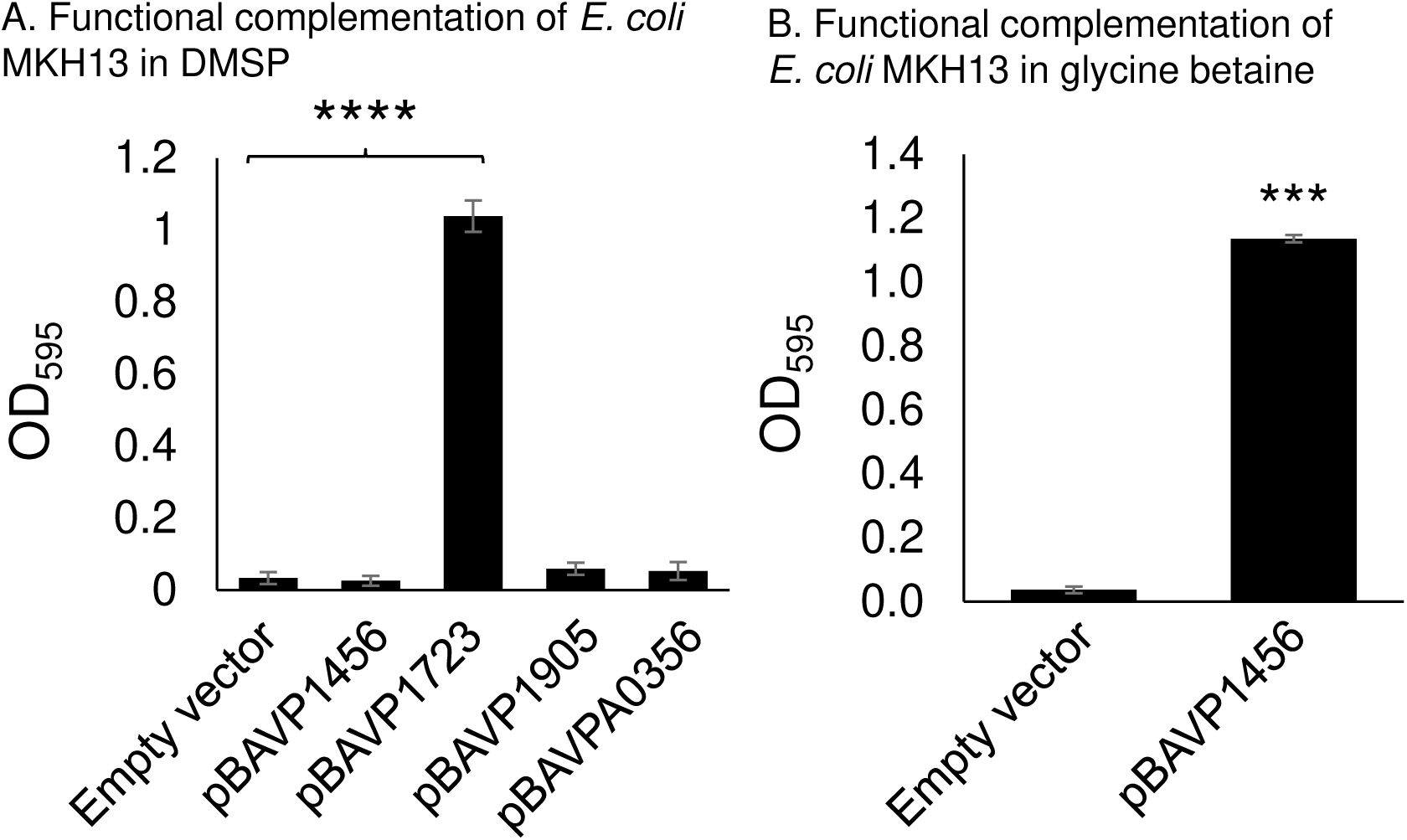
*E. coli* strain MKH13, which has deletions in all compatible solute transporters, was grown in M9G supplemented with 4% NaCl and functionally complemented with *V. parahaemolyticus* RIMD2210633 **(A)** *bccT1, bccT2, bccT3*, or *bccT4*, each expressed from an arabinose-inducible expression plasmid, pBAD33. Strains were grown for 24 hours in the presence of 500 µM DMSP and the final optical density (OD_595_) was compared to that of a strain harboring empty pBAD33. **(B)** *E. coli* MKH13 complemented with *bccT1* was grown in presence of 500 µM glycine betaine and compared to a strain harboring empty pBAD33. Mean and standard error of at least two biological replicates are shown. Statistics were calculated using a Student’s t-test (****, P < 0.0001).

### BccT3 from *V. cholerae* is a DMSP transporter

Of the four additional *Vibrio* species that used DMSP as an osmolyte, *V. harveyi* and *V. fluvialis* contain homologs of BccT1 and BccT2, which transport DMSP in *V. parahaemolyticus*. However, *V. vulnificus* contains homologs of only BccT3 and ProU2, while *V. cholerae* possesses only a BccT3 homolog and no ProU transporter. To examine whether BccT3 homologs from these species can uptake DMSP, we cloned *V. vulnificus bccT3* (VV2103, VVbccT3) and *V. cholerae bccT3* (FY484_RS06475, VCbccT3) homologs into pBAD33 and performed functional complementation assays in *E. coli* strain MKH13. We tested growth of MKH13 in the presence of glycine betaine to determine whether these transporters were functional, and found that both were glycine betaine carriers, as evidenced by growth of the MKH13 strain in M9G4% (**Fig. 6A**). Next, we examined DMSP uptake by these transporters and show that VCBccT3 functionally complemented MKH13, while VVBccT3 did not, similar to VpBccT3 (**Fig. 6B**).

**Figure 6.**
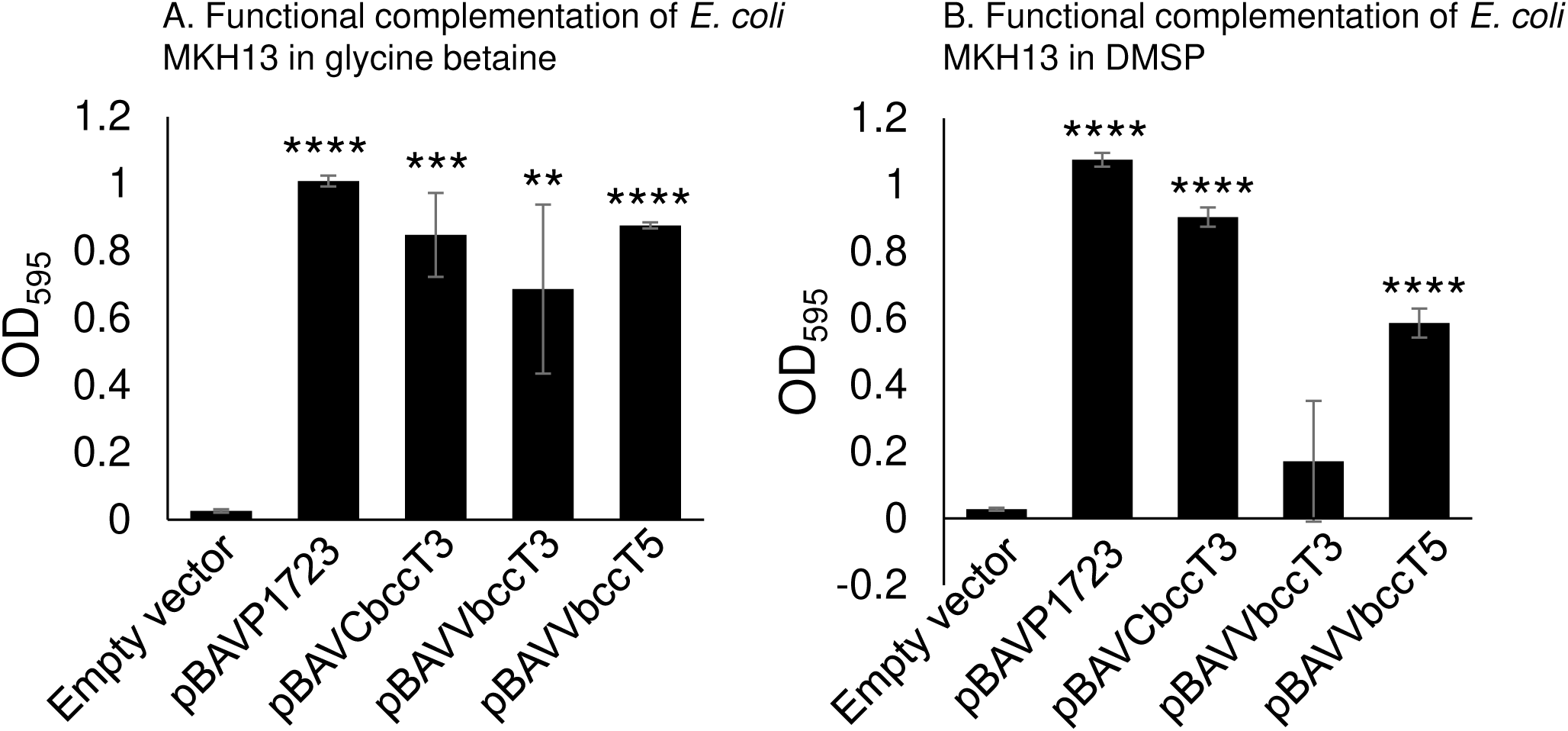
*E. coli* strain MKH13, which has deletions in all compatible solute transporters, was grown in M9G supplemented with 4% NaCl and functionally complemented with VP*bccT2* (pBAVP1723), VC*bccT3*, VV*bccT3* or VV*bccT5*, each expressed from an arabinose-inducible expression plasmid, pBAD33. Strains were grown for 24 hours in the presence of **(A)** 500 uM glycine betaine or **(B)** 500 µM DMSP and the final optical density (OD_595_) was compared to that of a strain harboring empty pBAD33. Mean and standard error of three biological replicates are shown. Statistics were calculated using a Student’s t-test (**, P < 0.01; ***, P < 0.001; ****, P < 0.0001).

### BccT5, an additional BCCT transporter present in *V. vulnificus*

Since BccT3 from *V. vulnificus* did not transport DMSP, we reexamined the *V. vulnificus* genome and identified an additional BCCT family transporter that shared < 32% homology with BCCTs from *V. parahaemolyticus*. This BCCT, which we named BccT5 (VV0783), was annotated as a 637-amino acid protein and was demonstrated via hydropathy profile analysis to have 12 TM domains (**Fig. S3**). This VVBccT5 had an 8 amino acid N-terminus and a 95 amino acid C-terminal tail. We complemented *E. coli* MKH13 with BccT5 expressed from pBAD33 and examined growth in M9G 4% NaCl supplemented with glycine betaine or DMSP. The *E. coli* construct was unable to grow in the presence of DMSP, and grew poorly in the presence of glycine betaine (data not shown). Further analysis of the nucleotide sequence of *bccT5* identified an alternative start site, which resulted in a larger 672-amino acid protein. This larger protein had a 43 amino acid N-terminus and 95 amino acid C-terminal tail. We cloned this protein into an expression plasmid and examined functional complementation of *E. coli* MKH13, which resulted in growth in the presence of either glycine betaine or DMSP (**Fig. 6**). This indicates that VVBccT5 can transport DMSP, along with VCBccT3. Thus, *Vibrio* species have at least four BCCT family transporters, including VPBccT1 and VPBccT2, that can uptake DMSP.

### Phylogenetic Analysis of BccTs among the Harveyi clade

VVBccT5 is a previously uncharacterized transporter that was undetected in a previous analysis. Using this protein as a seed, we examined members of the *Vibrionaceae* to determine BccT5’s distribution and evolutionary relationships. This analysis identified 53 *Vibrio* species, 15 *Photobacterium* species, 5 *Aliivibrio* species, two *Grimontia* species and 4 *Enterovibrio* species that contain a BccT with greater than 70% amino acid identity with 95-100% query coverage to VvBccT5 (**Fig. S4**). The branching pattern of BccT5 among these genera indicate a polyphyletic origin and that BccT5 is highly divergent within and among genera. The BccT5 tree contains two major lineages, I and II, with 100% bootstrap support. Within lineage I, three divergent clusters A, B, and C are present, with high bootstrap support, that are comprised of BccT5 from 43 *Vibrio* species. Within lineage II, 4 divergent clusters are present, D, E, F, and G, that are comprised of five genera, with high bootstrap values. BccT5 proteins from 11 *Photobacterium* species are present within cluster D, while cluster E branches distantly from D and is comprised of three *Vibrio* species. Within cluster F are BccT5 proteins from five *Aliivibrio* species that cluster closely with BccT5 proteins from *Shewanella* species, which are not members of the *Vibrionaceae* but are marine species. Within cluster G are six *Vibrio* species that branch together and on three additional separate branches are species belonging to *Grimontia, Enterovibrio* and four species of *Photobacterium* that cluster with *Enterovibrio* species. Overall, the clustering and branching pattern of BccT5 indicates that this protein does not have one origin and is a divergent protein that has been acquired several times within the *Vibrionaceae*.

Next, we wanted to determine the diversity of BccT proteins present among members of the Harveyi clade since in the BccT5 tree nearly all members of this clade, with the exception of *V. parahaemolyticus*, contained this protein. This analysis revealed the presence of at least 11 divergent BccT proteins within the Harveyi clade, with 100% bootstrap values and we named these BccT1 to BccT11 (**Fig. 7**). The majority of *V. parahaemolyticus* strains had four BCCT transporters, BccT1, BccT2, BccT3 and BccT4, with most other species having 7 BCCTs, *V. owensii* and *V. diabolicus* having 8 BCCTs, and *V. jasicida* having 9 BCCTs. In *V. jasicida* strains whose genome sequence is available, five BCCTs were present on chromosome 1 ranging in size from 523-632 amino acids and four BCCTs were present on chromosome 2 ranging in size from 511 to 535 amino acids. Overall, the tree demonstrates that there is significant diversity among BccTs from the Harveyi clade and even within clusters there is a great deal of protein divergence that could suggest functional divergence. An example of this is the BccT3 cluster, which shows members of the Cholerae clade, *V. cholerae, V. mimicus and V. metoecus* branching divergently from other members, and our growth analysis data showed that VCBccT3 can uptake DMSP but VVBccT3 and VPBccT3 cannot.

**Fig. 7.**
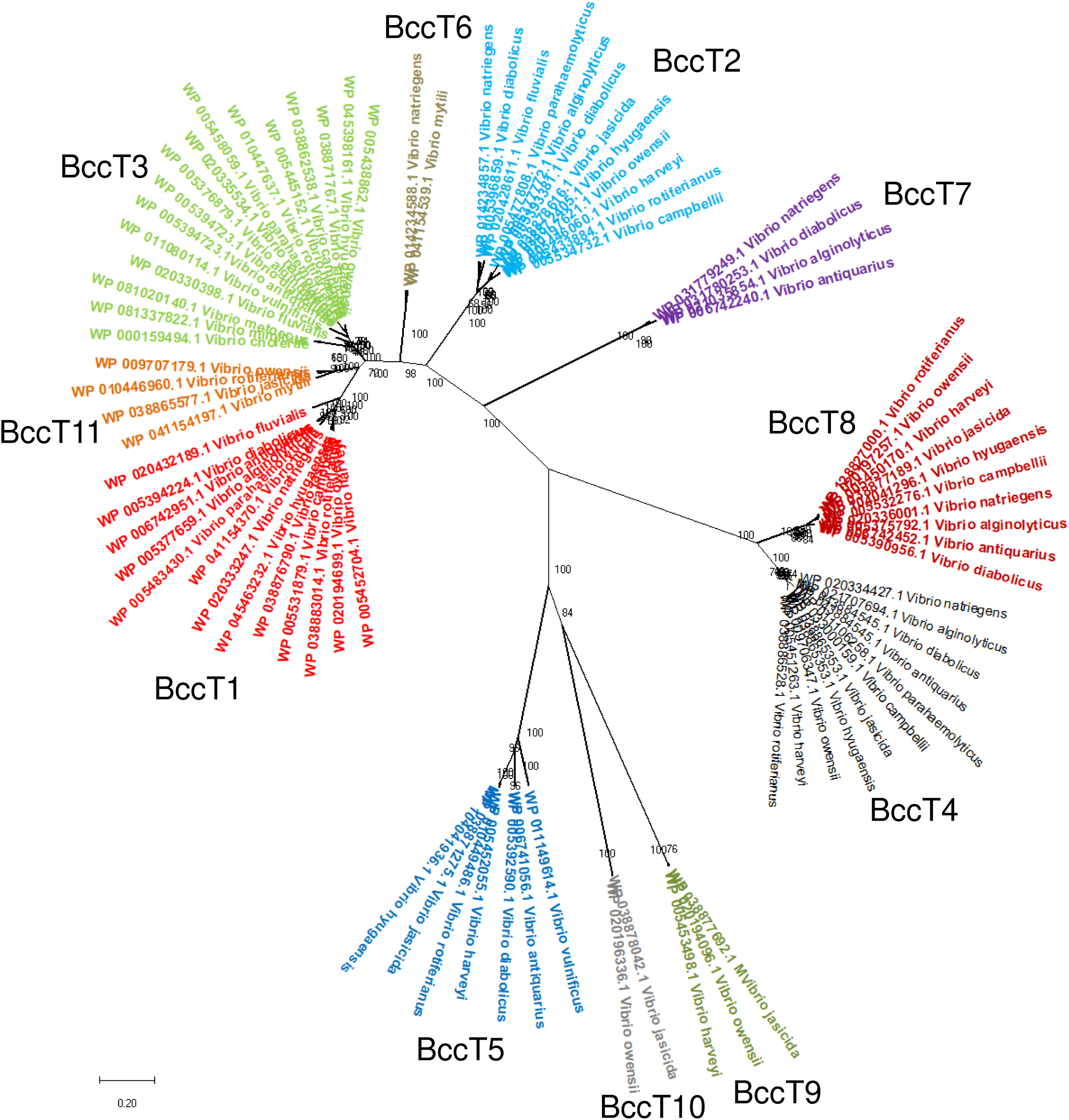
Phylogenetic tree of BccTs from the Harveyi clades and *V. cholerae, V. vulnificus and V. fluvialis* strains. A neighbor joining tree is shown with the sum of branch length = 9.31404349 and bootstrap values shown on each branch.

## Discussion

Here, we have shown that marine bacteria in the genus *Vibrio* can use DMSP as an effective osmolyte. These are the first studies to demonstrate a more widespread use of DMSP among marine halophiles. We demonstrate the important role BCCT carriers play in this process. BCCT-family transporters were solely responsible for the uptake of DMSP by *V. parahaemolyticus* and most likely BCCTs also play a role in DMSP uptake in other *Vibrio* species. Bacteria have adapted to hyper-osmotic conditions by changing the level of osmolytes in their cells to maintain turgor pressure. The most prevalent and energetically effective method to accomplish this is the uptake of osmolytes from the surrounding environment. DMSP is produced in vast quantities by marine phytoplankton as an osmoprotectant. This work showed that at least four different BCCT family proteins are carriers of DMSP in these marine species and these transporters are prevalent among this group.

Recently, an increased incidence of algal blooms has led to increased *V. parahaemolyticus* and *V. vulnificus* proliferation (76-78). Our data shows that *V. parahaemolyticus* does not utilize DMSP as a carbon source, but can grow and divide rapidly in high salinity when exogenous DMSP is present. We speculate that the ability of *V. parahaemolyticus* and other *Vibrio* species to utilize DMSP as a compatible solute increases their ability to proliferate in algal blooms that occur in both high salinity and in warmer months. Dinoflagellates are some of the main producers of DMSP and *V. parahaemolyticus* proliferation has been found in conjunction with dinoflagellate algal blooms (77, 78). Likewise, *V. vulnificus* can grow and divide rapidly and in higher salinity when exogenous DMSP is present.

Interestingly, our examination of the genome context of several BCCT family proteins identified associations with uncommon metabolic pathways in *Vibrio* species. For example, the *bccT6* genes that encodes BccT6 clustered with *gbcA* and *gbcB*, genes that are required for the catabolism of glycine betaine to dimethylglycine. In addition, this three-gene region is present in several additional species outside of the Harveyi clade suggesting that *Vibrio* species can catabolize glycine betaine (data not shown). The *bccT5* gene from *V. diabolicus* and *V. antiquarius* that encodes BccT5, clusters with a three gene operon *aldCalsSadhC2* coding for an alpha-acetolactate decarboxylase, alpha-acetolactate synthase (alpha-ALS), and acetoin oxidoreductase, respectively. In *Klebsiella* and *Enterobacter* species, these enzymes are required for the biosynthesis of 2,3-butanediol from pyruvate, a pathway that has not been previously identified in *Vibrio* species. This operon is also present in several additional *Vibrio* species including *V. alginolyticus* and *V. anguillarum*, but in these species, the region is associated with an ABC-type transporter. In the four species of *Vibrio* that contain BccT7, the *bccT7* gene clusters with a four-gene operon required for the catabolism of aromatic compounds and consists of genes encoding a glycine oxidase, a rubredoxin, an acyl-CoA reductase and an hydroxylating oxygenase. Although these examples are all association, they do suggest possible roles beyond osmolyte uptake for BccTs in *Vibrio* species.

## Methods

### Bacterial strains, media and culture conditions

All strains and plasmids used in this study are listed in Table 1. A streptomycin-resistant *V. parahaemolyticus* RIMD2210633 was used as the wild-type (WT) strain. *Vibrio parahaemolyticus* were grown either in lysogeny broth (LB) (Fisher Scientific, Fair Lawn, NJ) with 3% (wt/vol) NaCl (LB3%) or M9 minimal media (47.8 mM Na_2_HPO_4_, 22mM KH_2_PO_4_, 18.7 mM NH_4_Cl, 8.6 mM NaCl; Sigma-Aldrich) supplemented with 2mM MgSO_4_, 0.1 mM CaCl_2_, 20 mM glucose as the sole carbon source (M9G) and NaCl (wt/vol), as indicated. Dimethylsulfoniopropionate (DMSP) was used at a final concentration of 20 mM when supplied as a carbon source. *E. coli* strains were grown either in LB supplemented with 1% NaCl (LB1%) or M9G supplemented with 1% NaCl (M9G1%). All strains were grown at 37 C with aeration. Antibiotics were used at the following concentrations, as necessary: chloramphenicol (Cm), 25 µg/mL.

**Table 1.** Strains and plasmids used in this study.

| Strain | Genotype or description | Reference or Source |
| --- | --- | --- |
| *Vibrio parahaemolyticus* |  |  |
| RIMD2210633 | O3:K6 clinical isolate, Str<sup>r</sup> | (15, 86) |
| $\Delta ectB$ | RIMD2210633 $\Delta ectB$ (VP1721), Str<sup>r</sup> | (33) |
| SOYBCCT124 | RIMD2210633 $\Delta VP1456 \Delta VP1723 \Delta VPA0356$, Str<sup>r</sup> | (35) |
| SOYBCCT123 | RIMD2210633 $\Delta VP1456 \Delta VP1723 \Delta VP1905$, Str<sup>r</sup> | (35) |
| SOYBCCT134 | RIMD2210633 $\Delta VP1456 \Delta VP1905 \Delta VPA0356$, Str<sup>r</sup> | (35) |
| SOYBCCT234 | RIMD2210633 $\Delta VP1723 \Delta VP1905 \Delta VPA0356$, Str<sup>r</sup> | (35) |
| SOYBCCT1342 | RIMD2210633 $\Delta VP1456 \Delta VP1723 \Delta VP1905 \Delta VPA0356$ Str<sup>r</sup> | (35) |
| $\Delta proU1$ | RIMD2210633 $\Delta proU1$, Str<sup>r</sup> | (33) |
| $\Delta proU1 \Delta proU2$ | RIMD2210633 $\Delta proU1 \Delta proU2$, Str<sup>r</sup> | This study |
| *Vibrio harveyi* 393 | Isolated from barramundi in Australia | (87) |
| *Vibrio fluvialis* ATCC33809 | [NCTC 11327][606] Clinical isolate | (88) |
| *Vibrio vulnificus* YJ016 | Clinical isolate | (89) |
| *Vibrio cholerae* N16961 | O1, El Tor strain, Bangladesh, Clinical, 1975 | (90) |
| *Escherichia coli* |  |  |
| DH5α λpir | $\Delta lac$ *pir* | ThermoFisher Scientific |
| β2155 λpir | $\Delta dapA::erm$ *pir* for bacterial conjugation | (91) |
| MKH13 | MC4100 ($\Delta betTIBA$) $\Delta (putPA)101$<br>$\Delta (proP)2 \Delta (proU)$; Sp<sup>r</sup> | (75) |
| **Plasmids** |  |  |
| pBAD33 | Expression vector; *araBAD* promoter; Cm<sup>r</sup>; p15a origin | (80) |
| pBAVP1456 | pBAD33 harboring full-length VP1456 (*bccT1*) | (36) |
| pBAVP1723 | pBAD33 harboring full-length VP1723 (*bccT2*) | (35) |
| pBAVP1905 | pBAD33 harboring full-length VP1905 (*bccT3*) | (35) |
| pBAVPA0356 | pBAD33 harboring full-length VPA0356 (*bccT4*) | (36) |
| pBAVCBccT3 | pBAD33 harboring full-length BccT3 from *V. cholerae* | This study |
| pBAVVBccT3 | pBAD33 harboring full-length BccT3 from *V. vulnificus* | This study |
| pBAVCBccT5 | pBAD33 harboring 637 amino acid BccT5 from *V. vulnificus* | This study |
| pBAVCBccT5long | pBAD33 harboring 672 amino acid BccT5 from *V. vulnificus* | This study |

### Mutant strain construction

The Δ*proU1*Δ*proU2* double mutant was constructed by creating an in-frame deletion of *proU2* in a *proU1* background (33). Splicing by overlap extension PCR and allelic exchange was performed as previously described (79). *V. parahaemolyticus* RIMD2210633 genomic DNA template and SOE primers listed in Table 2 were used in the SOE PCR amplification to create a truncated *proU2* PCR fragment of 750 bp in size by removing ∼1,150 bp from the wild-type *proU2* gene (1,900 bp). The truncated *proU2* fragment was subsequently cloned into a pDS132 suicide vector, transformed in *E. coli* β2155 λpir DAP auxotroph, and mobilized into a recipient wild type *V. parahaemolyticus* RIMD2210633 strain by conjugation (33). The generated *V. parahaemolyticus* RIMD2210633 mutant strain harboring a truncated version of the *proU2* gene was designated Δ*proU2*. To create the double mutant Δ*proU1*Δ*proU2* strain, *V. parahaemolyticus* RIMD2210633 Δ*proU2* strain was used as background, and a previously created Δ*proU1* inserted into the Δ*proU2* background by conjugation and homologous recombination. All deletions were confirmed by PCR and sequencing.

**Table 2.**
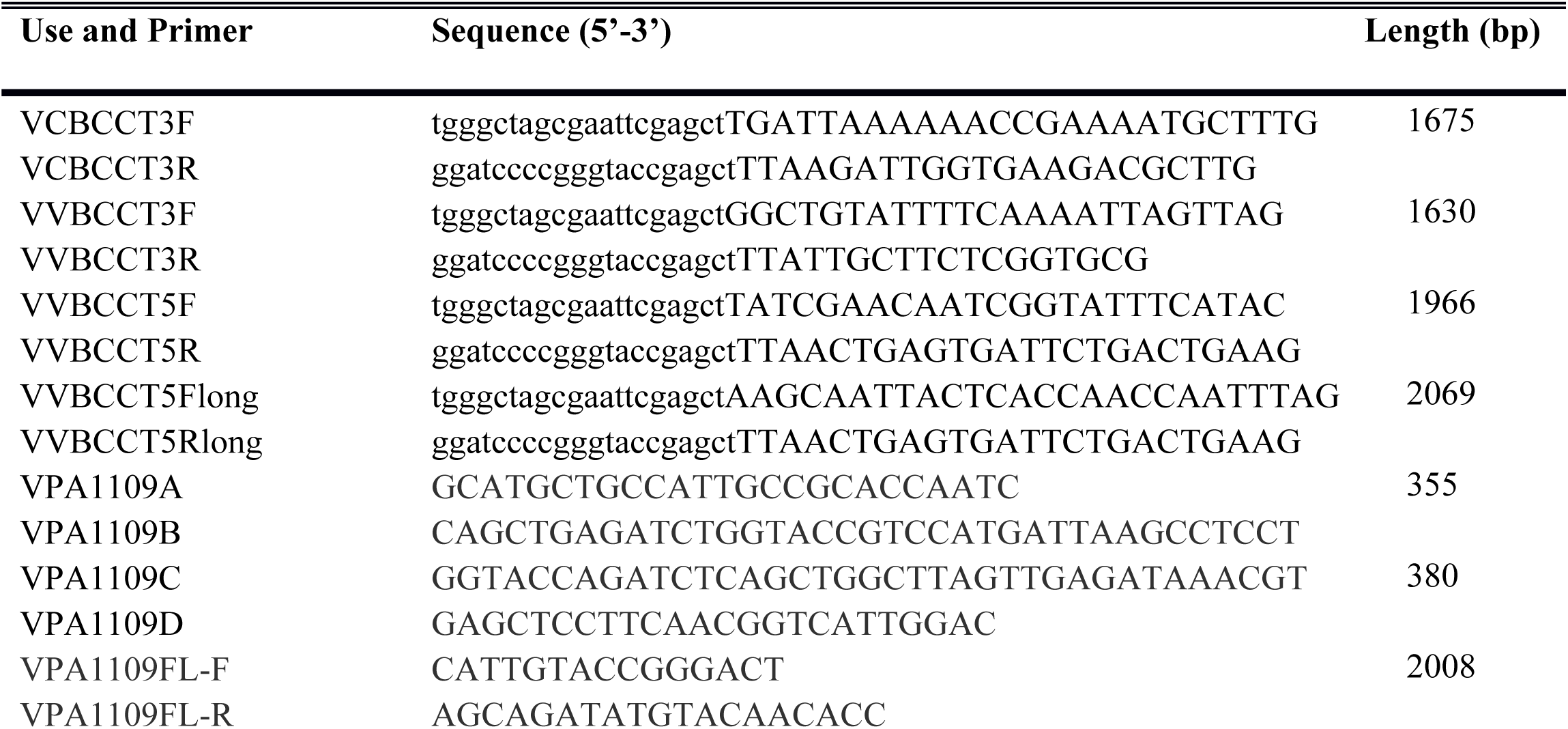
Primers used in this study.

### Growth analysis of mutant strains in DMSP

*V. parahaemolyticus* RIMD2210633 or an in-frame deletion mutant of *ectB* were grown overnight in M9G1%. Cultures were subsequently diluted 1:50 into fresh medium and grown for five hours and used as previously described for analysis of DMG as an osmolyte (36). These strains and the *proU* double mutant, and a quadruple Δ*bccT1*Δ*bccT3*Δ*bccT4*Δ*bccT2* mutant (*bccT* null) strains exponential cultures were then diluted 1:40 into 200 µL of M9G6% medium with and without exogenous compatible solutes in a 96-well microplate and grown at 37° C with intermittent shaking for 24 hours. Dimethylsulfoniopropionate (DMSP) was added to a final concentration of 500 µM. Growth analysis was repeated following the above procedure with each of four triple *bccT* deletion mutants, Δ*bccT2*Δ*bccT3*Δ*bccT4*, Δ*bccT1*Δ*bccT3*Δ*bccT4*, Δ*bccT1*Δ*bccT2*Δ*bccT4*, Δ*bccT1*Δ*bccT2*Δ*bccT3*, to determine which BCCT was responsible for transport of DMSP.

### Functional complementation of *E. coli* strain MKH13

The 1623-bp *opuD* (*bccT3* homolog) was amplified from the *V. cholerae* N16961 genome using the primers listed in Table 2. The 1572-bp bccT3 homolog, the 1914-bp *bccT5* or the 2016-bp *bccT5* gene were amplified from the *V. vulnificus* YJ016 genome using primers listed in Table 2. All primers were purchased from Integrated DNA Technologies (Coralville, IA). Gibson assembly protocol using NEBuilder HiFi DNA Assembly Master Mix (New England Biolabs, Ipswich, MA) was followed to ligate the amplified fragments with the expression vector pBAD33 (80), which had been linearized with SacI. Regions of complementarity for Gibson assembly are indicated by lowercase letters in the primer sequence in Table 2. The resulting expression plasmids, pBAVCbccT3, pBAVVbccT3, pBAVVbccT5, or pBAVVbccT5long, were transformed into *E. coli* Dh5α for propagation. Plasmids were then purified, sequenced, and subsequently transformed into *E. coli* strain MKH13, which has large deletions that include compatible solute transporters (*putP, proP, proU*) and the choline uptake and GB biosynthesis loci (*betT*-*betIBA*) (75).

*E. coli* MKH13 strains containing pBAD expressing a single *bccT* gene were grown overnight in minimal media supplemented with 1% NaCl and 20 mM glucose (M9G1%) with chloramphenicol and subsequently diluted 1:100 into M9G supplemented with 4% NaCl (M9G4%) and 500 µM of the indicated compatible solute and chloramphenicol. Expression of each BCCT was induced with 0.01% arabinose and functional complementation was determined by measuring OD_595_ after 24 hours growth at 37°C with aeration. Growth was compared to that of an MKH13 strain harboring empty pBAD33, which cannot grow in M9G4% without exogenous compatible solutes. Statistics were calculated using a Student’s t-test.

### Bioinformatics and phylogenetic analyses

Transmembrane helix probabilities of *V. parahaemolyticus* BccT1 (Q87PP5.1) and *V. vulnificus* BccT5 (BAC93547.1) were generated using OCTOPUS and aligned via the AlignMe program (http://www.bioinfo.mpg.de/AlignMe) (81-83). A Neighbor-Joining tree for BccTs among the Harveyi clade was constructed using MEGAX (84). The evolutionary distances were computed using the Jones-Taylor_thornton (JTT) matrix-based method and are in the units of the number of amino acid substitutions per site. The rate variation among sites was modeled with a gamma distribution (shape parameter = 5). This analysis involved 84 amino acid sequences. There were a total of 737 positions in the final dataset. A phylogenetic tree for BccT5 was constructed using the Maximum Likelihood method and Le_Gascuel_2008 model conducted in MEGA X (84, 85). The percentage of trees in which the associated taxa clustered together is shown next to the branches. Initial tree(s) for the heuristic search were obtained automatically by applying Neighbor-Join and BioNJ algorithms to a matrix of pairwise distances estimated using a JTT model, and then selecting the topology with superior log likelihood value. A discrete Gamma distribution was used to model evolutionary rate differences among sites (3 categories. The trees are drawn to scale, with branch lengths measured in the number of substitutions per site. For BccT5 tree, 88 amino acid sequences with a total of 623 positions were examined. All positions with less than 95% site coverage were eliminated, i.e., fewer than 5% alignment gaps, missing data, and ambiguous bases were allowed at any position (partial deletion option).

## ACKNOWLEDGEMENTS

This research was supported by a National Science Foundation grant (award IOS-1656688) to E.F.B. G.J.G. was funded in part by a University of Delaware graduate fellowship award. We thank members of the Boyd Group for constructive feedback on the manuscript.

---

**Figure S1**. Growth analyses of *V. parahaemolyticus* RIMD2210633 (down triangles), *V. harveyi* 393 (up triangles), *V. vulnificus* YJ016 (diamonds), *V. cholerae* N16961 (squares), and *V. fluvialis* (circles), in M9 with DMSP (open shapes) as the sole carbon source or M9G (solid shapes). Optical density (OD_595_) was measured every hour for 24 hours. Mean and standard error of two biological replicates are shown.

**Figure S2**. Growth analysis of wild-type and *proU1proU2* double mutant was conducted in M9G supplemented with 6% NaCl and DMSP. Optical density (OD_595_) was measured every hour for 24 hours; mean and standard error of at least two biological replicates are displayed.

**Figure S3**. The transmembrane helix probability of *V. vulnificus* YJ016 BccT5 was generated and aligned with *V. parahaemolyticus* RIMD2210633 BccT1 using AlignMe (http://www.bioinfo.mpg.de/AlignMe). Values close to 1 indicate a high probability of that sequence being in the membrane while 0 is a low probability of that sequence being in the membrane. Dots below the plot indicate gaps introduced during alignment.

**Figure S4**. Maximum Likelihood tree with the highest log likelihood (−14924.55) is shown for BccT5. Bootstrap values are shown next to each branch. Two main lineages are shown, I subdivided into three groups A, B, and C and linage II subdivided into 4 main groups. Species with the same genus are colored identically.

